# spRefine Denoises and Imputes Spatial Transcriptomics with a Reference-Free Framework Powered by Genomic Language Model

**DOI:** 10.1101/2025.04.22.649977

**Authors:** Tianyu Liu, Tinglin Huang, Wengong Jin, Tinyi Chu, Rex Ying, Hongyu Zhao

## Abstract

The analysis of spatial transcriptomics is hindered by high noise levels and missing gene measurements, challenges that are further compounded by the higher cost of spatial data compared to traditional single-cell data. To overcome this challenge, we introduce **spRefine**, a deep learning framework that leverages genomic language models to jointly denoise and impute spatial transcriptomic data. Our results demonstrate that spRefine yields more robust cell- and spot-level representations after denoising and imputation, substantially improving data integration. In addition, spRefine serves as a strong framework for model pre-training and the discovery of novel biological signals, as highlighted by multiple downstream applications across datasets of varying scales. Notably, spRefine enhances the accuracy of spatial ageing clock estimations and uncovers new aging-related relationships associated with key biological processes, such as neuronal function loss, which offers new insights for analyzing ageing effect with spatial transcriptomics.

## 1 Introduction

Spatially-resolved transcriptomic (SRT) technologies have enabled the investigation of the cellular functions under the spatial context [1], which cannot be directly accessed through single-cell RNA sequencing (scRNA-seq) technology. SRT technologies fall into two primary categories: (1) Imaging-based methods, including smFISH [2, 3], MERFISH [4], seqFISH [5] and Xenium [6, 7]; and (2) Sequencing-based methods, including Visium, STARmap [8] and Slide-seq [9]. Imaging-based methods can measure gene expression at the subcellular resolution and generate single-cell gene expression profiles through aggregation. However, the small area covered as well as the limited number of genes pose challenges in characterizing the spatial context of the target tissue comprehensively [10, 11]. On the other hand, although sequencing-based methods can profile at the genome-wide level, they tend to have lower resolution and higher noise [12]. These challenges presented in these datasets hidden biological signals such as ageing-associated or survival-associated features [13]. Therefore, there is a need to develop computational tools to address the limitations of different platforms and improve the reliability of the interpretation of spatial transcriptomics.

Efforts have been made to address the limitations of these two platforms. For imaging-based technologies, researchers utilize reference scRNA-seq datasets with a larger number of measured genes to impute the unmeasured genes in the spatial transcriptomics, whose related methods were already benchmarked [14]. As examples, methods have been proposed to predict unmeasured gene expression by aggregating nearest neighbors of cells for regression [15–18], joint probabilistic modeling [19–22], and optimal transport [11, 23, 24]. However, imputation results are limited by the quality of scRNA-seq data, which may lead to challenges in interpreting the imputed profiles. For transcriptomics from sequencing-based methods, Sprod [25] was developed to integrate both image features and gene expression levels to improve data quality by graph smoothing. SEDR [12] improves data quality and generates better representations through an unsupervised learning framework. However, these methods are restricted to sequencing-based methods and cannot uncover the missing information from unmeasured genes, and they are not able to process large-scale SRT datasets. Therefore, it is critical to have a flexible and efficient method for processing datasets from different platforms as well for driving new biology discoveries.

Recently, a method has been proposed for imputing imaging-based SRT datasets from gene networks based on protein language models (PLMs) [26], which play an important role in the unmeasured gene expression prediction task according to their ablation tests. PLMs are foundation models trained with large-scale amino-acid sequences to model protein data as a language [27, 28], and the success of applying PLMs to this task suggests the utility of PLMs to build expression relationships among genes. However, this method is limited to the imputation of protein-coding genes. To simultaneously impute and denoise gene expression profiles, we may consider Genomic Language Models (gLMs) [29–31], which model DNA sequence as a language, to derive reliable gene-gene correlations in the expression levels. There exist gLMs pre-trained with either DNA sequence information purely or jointly between DNA sequences and expression profiles, and the latter setting is more related to our problem. Furthermore, the imputation process does not require external information such as scRNA-seq data, which will be more flexible for SRT data without paired scRNA-seq data or SRT data on a large scale. In principle, leveraging gene-gene correlations from gLMs pre-trained with different gene expression profiles will also improve data quality.

In this manuscript, we present spRefine, a novel method to refine the quality of SRT data collected from different platforms. spRefine utilizes both measured gene expression profiles as well as gene embeddings from gLMs as inputs and learns the hidden gene-gene interactions in SRT data with a coupled Auto-Encoder, which generates spot embeddings and dataset-specific gene embeddings simultaneously. It then imputes and denoises the given data by multiplying the learned embeddings to output a new matrix with improved expression levels. We demonstrate better performance of spRefine through its application to SRT data of different scales from various platforms. With a larger number of inferred genes as well as better data quality, spRefine can reduce batch effect, discover novel cell-cell communications, and improve spatial ageing clock modeling. Moreover, spRefine can also work as a pre-training framework to unify cell and spot representations. In addition, we show that spRefine can leverage the disease-associated marker genes selected based on the imputed HEST-1k data [32] to better predict survival or disease states of patients from The Cancer Genome Atlas (TCGA) database [33]. Our results show the strong capacity of spRefine in transferring the learned knowledge to biomedical data with different resolutions and improving our understanding of ageing and cancer processes.

## 2 Results

### Overview of spRefine

We design spRefine with a decomposed auto-encoder for both imputing gene expression and denoising the measured gene expression of spatial transcriptomics. spRefine contains two modules for learning cell embeddings and gene embeddings separately, while the cell embeddings are generated based on measured expression profiles, and the gene embeddings are generated based on a pre-trained sequence-to-function model. To generate the final imputed data matrix, we multiply the cell embeddings and gene embeddings optimized by reconstructing the denoised expression profiles, shown in Figure 1 (a). Therefore, spRefine is capable of performing dataset-specific analysis for various downstream applications including spatial domain identification, cell-cell communication inference, batch effect correction, and spatial ageing clock improvement, with the help of task-specific tools shown in Figure 1 (b). spRefine can also work as a pre-training architecture, which treats imputation+denoising as a pre-training task and learns cell embeddings from large-scale spatial transcriptomics. spRefine performs better than baselines and can facilitate several downstream analyses with pre-training design, including survival prediction, phenotype identification, and disease-state diagnosis, shown in Figure 1 (c). Details of model development are discussed in the Methods section.

**Fig. 1.**
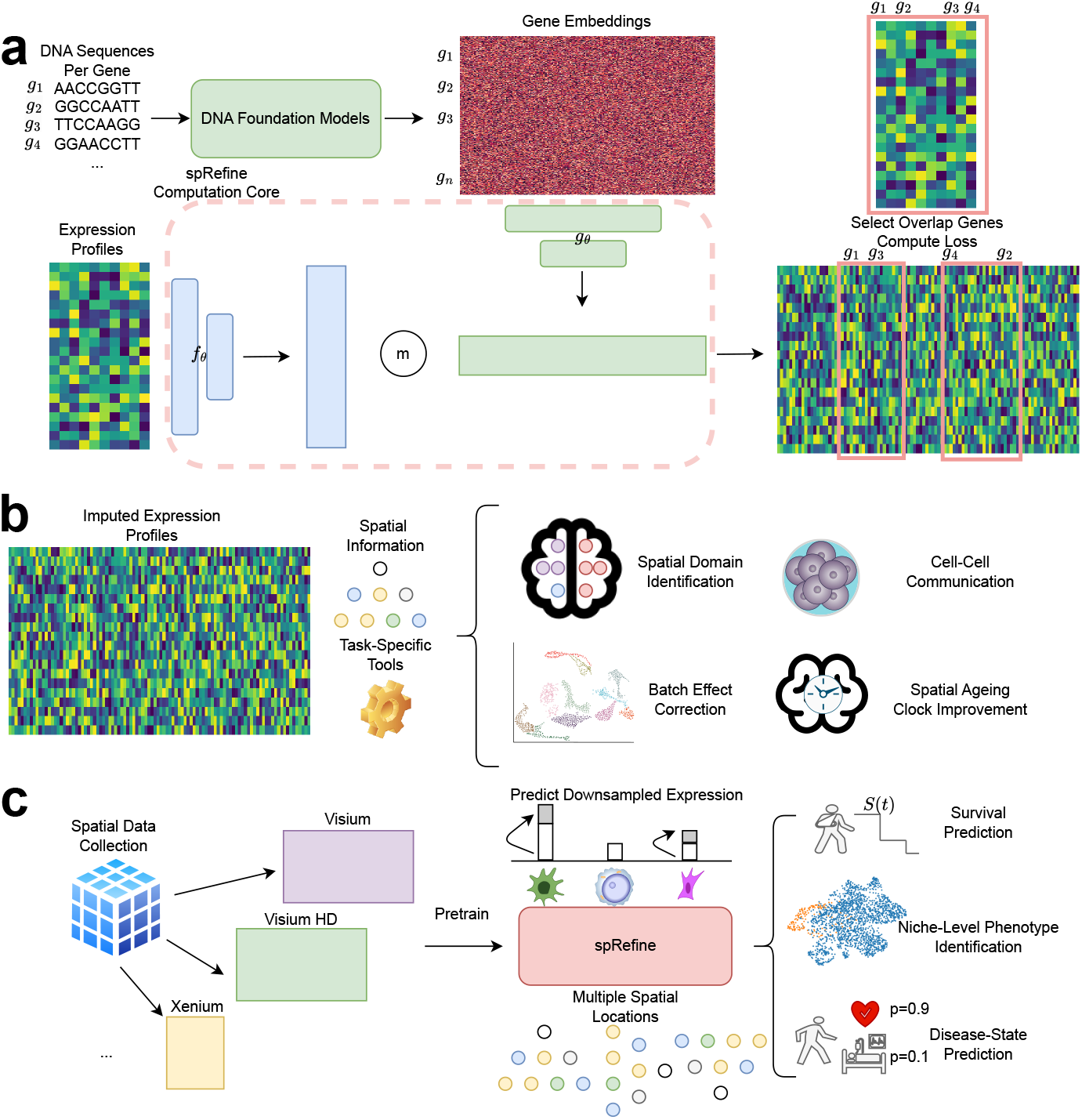
The landscape of spRefine and related applications. (a) The computation framework of spRefine as a tool for imputation and denoising. spRefine utilizes prior information from DNA foundation models to help uncover missing gene-gene interactions and gene expressions unmeasured by the raw transcriptomic profiles. (b) The applications of spRefine including spatial domain identification, cell-cell communication inference, batch effect correction, and spatial ageing clock improvement. (c) spRefine as a pre-training framework for representing phenotype information. spRefine is capable of performing survival prediction, identifying spot-level phenotype information, and predicting disease states.

### spRefine improves data quality through imputing profiles

spRefine performs imputation and denoising by predicting the unmeasured genes’ expressions and improving the observed ones. We call this process as “refine” in this manuscript. Inspired by [34], we designed a biology-driven framework for evaluating the refined gene expression profiles by assessing the discovery of biological signals by comparing our method with different baselines. By collecting observed cell-type labels, we tested the clustering performances based on the expression profiles and utilized Normalized Mutual Information (NMI), Adjusted Rand Index (ARI), Average Silhouette Width (ASW), and the averaged score (Avg) as metrics for the evaluation of signal discovery [35, 36]. Furthermore, we also considered demonstrating the contribution of refined profiles for correcting batch effect in the spatial transcriptomics, which can be evaluated by the metrics focusing on biological signal preservation (*S*_*bio*_) and batch effect removal (*S*_*batch*_) from scIB [36]. These metrics include Isolated Label Score, NMI, ARI, cell-type ASW (cASW), cell-type Local Inverse Simpson’s Index (cLISI), batch LISI (bLISI), batch ASW (bASW), k-nearest-neighbor Batch-Effect Test (kBET), Graph Connectivity Score (GC), and PCR comparison score. All of the metrics are in (0,1) and higher values mean better model performance. Details of our evaluation framework are summarized in the Methods section.

To measure the effectiveness of imputation, we considered some existing imputation methods, including VISTA [22], ENVI [20], Tangram [11], gimVI [19], TransImp [37], and SpaGE [17]. Most of these methods were found to have relatively good performance in benchmark studies [7, 14]. Moreover, to test if incorporating spatial information can improve cell-type annotation, we also aggregated the spots into niches with major-voting labels as a baseline for spatial effect (named as Niches). Furthermore, we included a foundation model trained with spatial transcriptomics, known as Novae [38], in our baseline to test if our imputed profiles can generate better representations than those pre-trained models. We considered four datasets with subcellular spatial transcriptomics (SST), named as xenium breast [39], xenium brain [39], seq-fish embryo [40], and osmfish brain [41]. According to Figures 2 (a)-(d), spRefine has the best overall performance across four datasets, followed by VISTA. Moreover, Novae and Niches performed worse than spRefine, suggesting that spRefine can better capture information than the pre-trained models or generated niches based on spatial location. Moreover, there was better cell type separation after imputation for the xenium breast sample, supported by the cell-type-level similarity change shown in Extended Data Figure 1 (a). spRefine is also robust to the change of random seeds, shown in Extended Data Figure 1 (b). We also performed several ablation tests, including the choices of initial gene embeddings, the choices of encoder, and the choices of loss function design, shown in Extended Data Figures 2 (a)-(c). Details are discussed in the Methods section. Finally, the details of hyper-parameter tuning can be found in the Methods section.

**Fig. 2.**
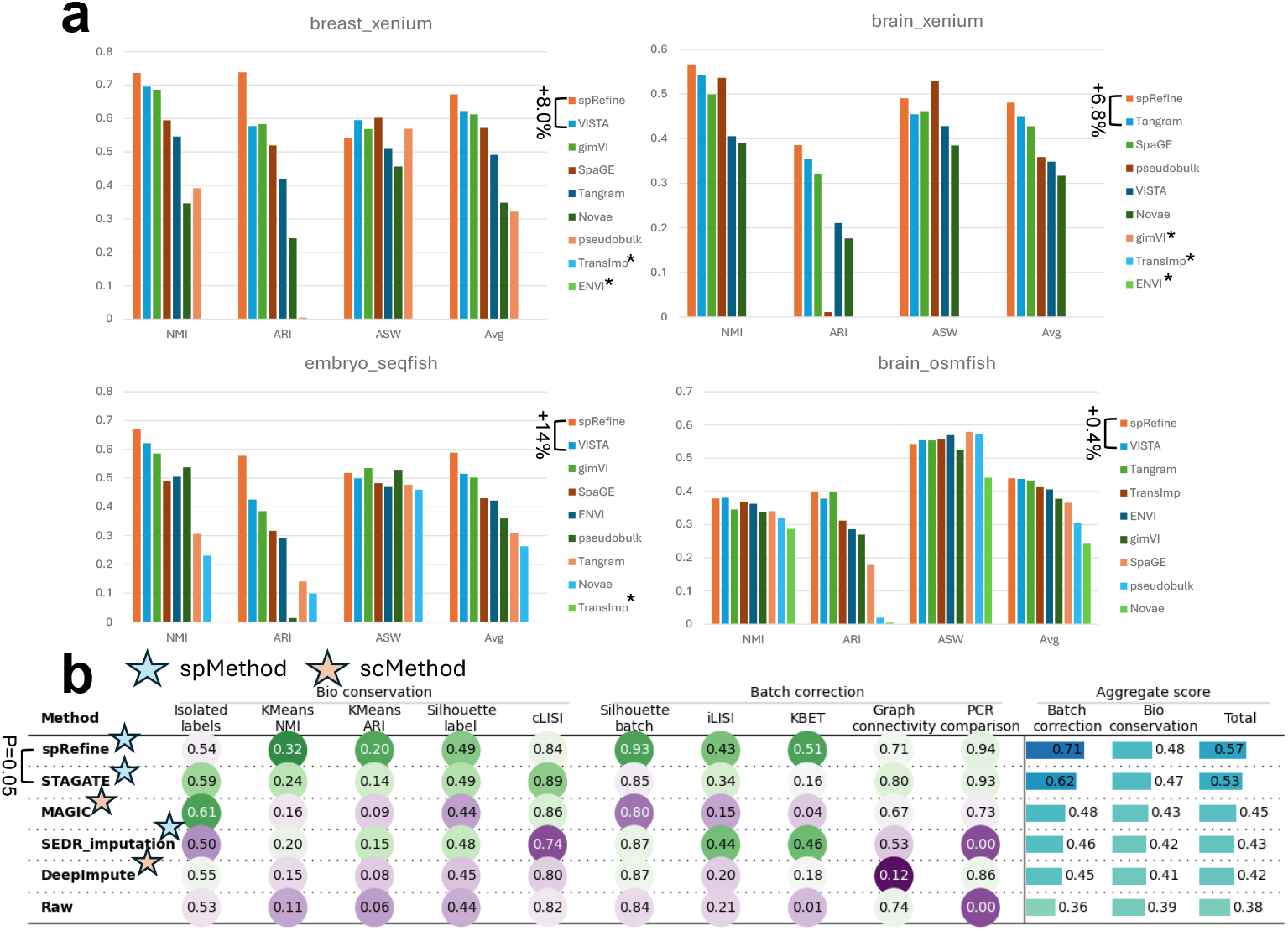
Comparisons between spRefine and other baselines for cell clustering and batch effect correction. (a) The clustering performance based on different imputation methods or spatial foundation models across four sub-cellular-resolved spatial transcriptomic data. The star means that the selected method met out-of-memory (OOM) errors in the specific dataset. The methods are ranked by the Avg score for each dataset, and we highlight the increment made by spRefine compared with the second-best baseline method. (b) The performance of batch effect correction based on different methods for processing the SpatialLIBD dataset. Here *spMethod* represents the method designed for spatial transcriptomics, while *scMethod* represents the method designed for single-cell transcriptomics. We performed the Wilcoxon Rank-sum test (one-side) between spRefine and the second-best baseline method, and annotated the p-value (P) in this panel.

To evaluate the performance of spRefine in denoising measured gene expression profiles, we followed the idea from [12], and utilized the batch effect correction task as an indicator to evaluate model performances in reducing the noise level of the given dataset. The baselines we considered include SEDR [12] and STAGATE [42], which were designed for spatial transcriptomics denoising and tested for reducing batch effect. We also considered representative denoising methods designed for single-cell transcriptomics, including MAGIC [43] and DeepImpute [44] according to their performances in benchmarking analysis [45]. We used the 10X Visium dataset from SpatialLIBD [46] for evaluation. This dataset contains multiple slides and also offers expert annotation of spot types to validate the preservation of biological signals. Our selected baselines provide recommended hyper-parameter settings based on this dataset for our evaluation. To perform a fair comparison, we imputed the SpatialLIBD dataset with different methods and utilized Harmony [47] for data integration. According to Figure 2 (e), spRefine outperformed the baselines in reducing the batch effect significantly, as reflected in *S*_*bio*_, *S*_*batch*_, and *S*_*total*_. Again, including a GNN-based encoder does not reduce the noise of tested spatial transcriptomic datasets, shown in Extended Data Figure 2 (d). We further visualized the Uniform Manifold Approximation and Projection (UMAP) [48, 49] before and after integration, shown in Extended Data Figure 3. This figure also demonstrates that spRefine with Harmony can help with data integration.

### Discovering novel cell-cell communications for breast cancer

Imputing spatial transcriptomics can facilitate cell-cell communication (CCC) inference in the spatial context [50] by expanding the list of genes. CCC is a technique used to capture the cell-cell interactions grouped by cell types from gene-gene interactions. Here we considered the breast cancer dataset and imputed gene expressions using spRefine. We compared the number of significant CCCs based on the raw and imputed profiles using COMMOT [51], which utilizes optimal transport [52] across spatial distributions to capture the dynamic process of gene expression changes and infer CCCs. According to Figure 3 (a), a higher number of CCCs (19 vs. 3) were inferred based on the imputed profiles, as expected. We further visualized the cell-type composition of the given sample in Figure 3 (b), which contains the location information of malignant cells as well as immune cells. It has been shown that chemokines play an important role in the development and migration of cancer cells, and thus we focused on the interaction of chemokines-related genes to understand the landscape of the microenvironment in breast cancer. According to Figures 3 (c) and (d), there exists a strong interaction between the sender and receiver at the junction of immune cells and ECM 1+/CRABP2 + Malignant cells, and the signal flow further validated this discovery. The role of CXCL12 and CXCR4 has been widely discussed in the mechanism and treatment of cancer, and thus this interaction is an important marker for microenvironment [53, 54]. Moreover, spRefine identified a novel CCC between CCL15 and CCR1, which was not inferred based on the raw dataset. According to Figures 3 (e) and (f), this CCC shows a strong interaction in the junction between immune cells and SCGB2A2+ malignant cells, as another key signal for breast cancer microenvironment [55, 56]. Therefore, spRefine can offer additional insights through more identified CCCs. The full list of cell-cell interactions inferred by spRefine is summarized in Extended Data Figures 4 and 5.

**Fig. 3.**
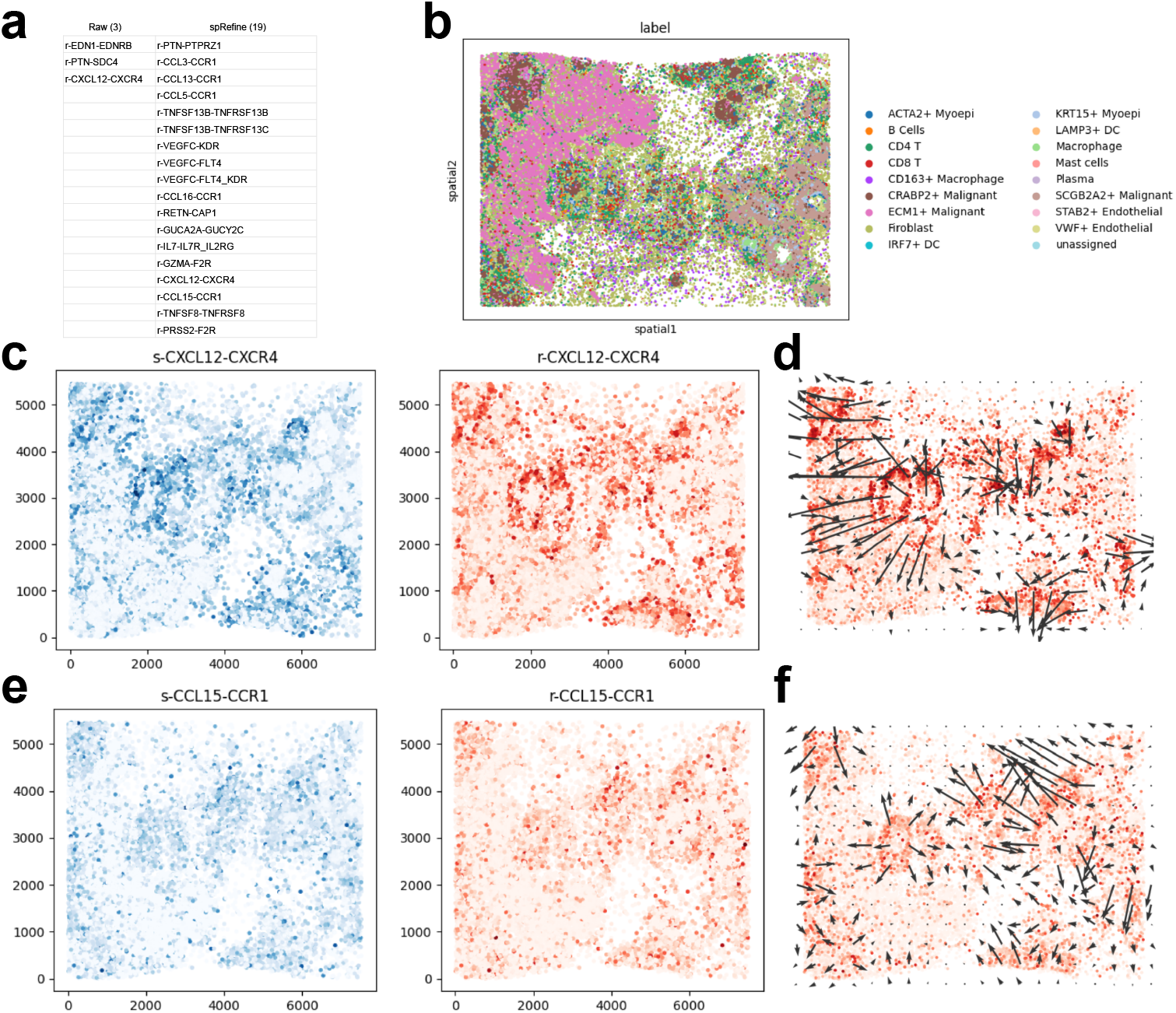
Using spRefine for discovering novel cell-cell interactions (CCIs). (a) The comparison of discovered signals between the results of COMMOT based on the raw expression profiles (Raw) and imputed expression profiles (spRefine). (b) The visualization of cells in xenium breast dataset colored by cell types based on spatial location. (c) The plots containing sender and receiver strength for the CCI CXCL12-CXCR4. (d) The signal direction of the CCI CXCL12-CXCR4. (e) The plots containing sender and receiver strength for the CCI CCL15-CCR1. (f) The signal direction of the CCI CCL15-CCR1.

### Utilizing imputed profiles as a pre-training framework for identifying disease-associated clusters

spRefine may work as an effective pre-training framework to preserve biological signals with the help of intrinsic gene-gene interaction supported by the embeddings from DNA foundation models. To validate this assumption, we collected large-scale spatial transcriptomics from two publicly available databases, including HEST-1k [32] and STimage-1K4M [57]. These two datasets contain data from different sequencing technologies with little overlap. There are more than 500 samples and some are annotated with spot identification, e.g., tumor cells versus normal cells from the patient sample. With such information, we could validate learned representations based on identifying the disease-level (or phenotype-level) information based on cell embeddings. Phenotype information (e.g., cancer cells) is usually treated as a different covariate compared with cell types [58], and thus we intend to examine if we can utilize pre-trained model to identify such covariates by clustering. The workflow in this section is summarized in Figure 4 (a).

**Fig. 4.**
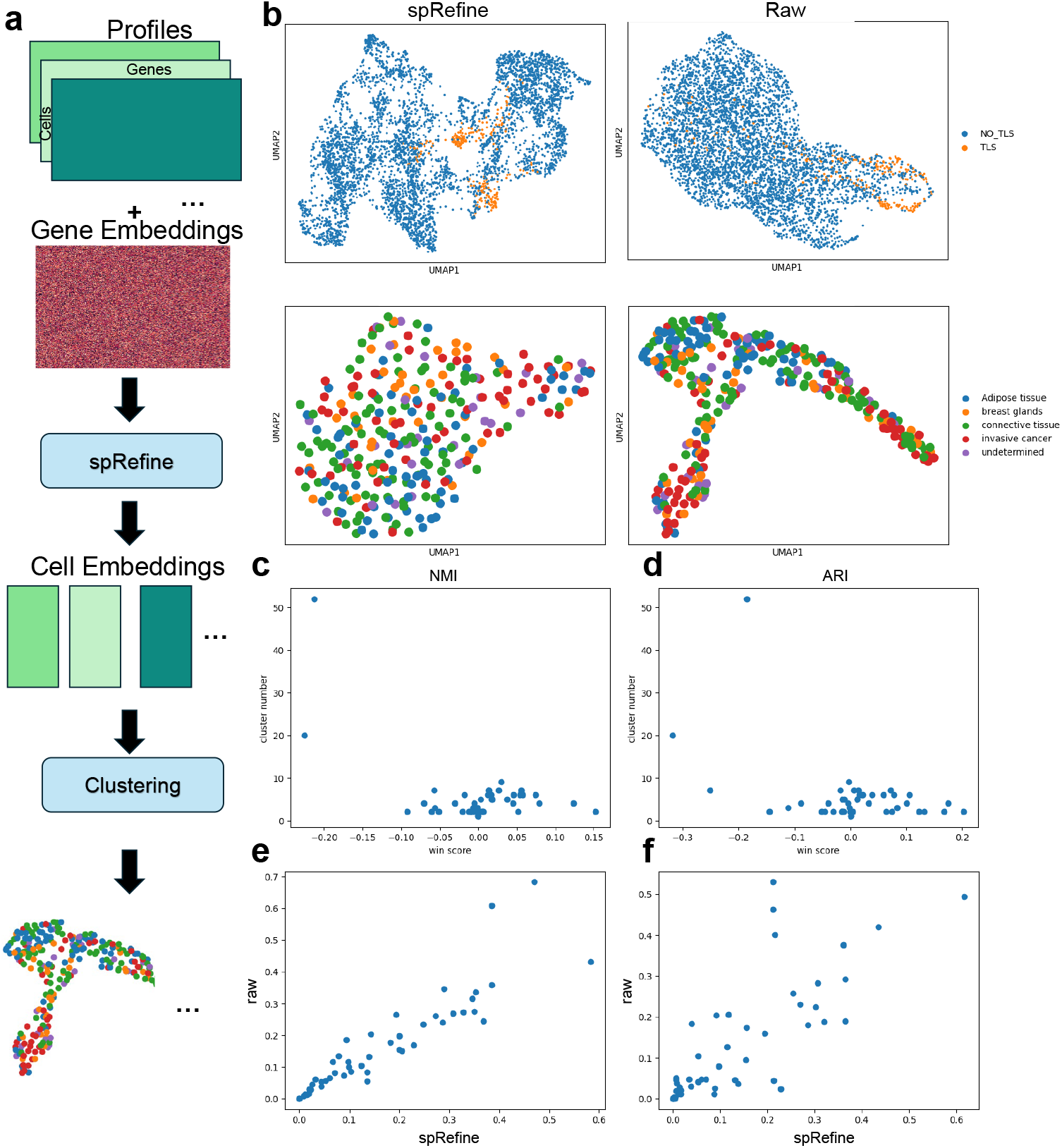
Analyzing the results of spRefine pre-trained with large-scale spatial transcriptomics. (a) The workflow of pre-training spRefine for phenotype-level identification. (b) The UMAPs for visually comparing spot representations before (right) and after (left) imputation based on the two selected samples with different cell states. (c) The relationship between win rate computed based on NMI and cluster number. (d) The relationship between win rate computed based on ARI and cluster number. (e) The relationship between the NMI scores of raw data and data imputed by spRefine. (f) The relationship between the ARI scores of raw data and data imputed by spRefine.

Here we pre-trained spRefine based on the combined datasets and using the expression enhancement and imputation approaches as the pre-training strategies. Different from our previous experiments, we focused on more diverse data and phenotype-level differences. We first visualize representative samples based on the UMAP colored by their cell-state labels, shown in Figure 4 (b). This figure contains samples whose clusters are either easy or difficult to identify after imputation. The easier sample contains cells with or without tertiary lymphoid structures (TLS), while the difficult sample contains a more complicated tissue structure, including five different annotations. We can see that imputation does not always identify specific cells or spots, and the more diverse the cells are, the more difficult it is to distinguish the phenotypes of cells by the profiles after imputation. We computed the clustering metrics NMI score and ARI score, as well as the win rate after imputation for these two scores, where win rate = *S*_*imputed*_ −*S*_*raw*_. The relationship between the number of the clusters in the tested samples and the win rates of two scores are presented in Figures 4 (c) and (d), which show a clear negative correlation between them (PCC=-0.55, p-value=3.91e5 for NMI, and PCC=-0,44, p-value=1.8e-3 for ARI). Therefore, imputation profiles are better suited to help identify cell types in samples with fewer predefined categories, for example, a sample with only tumor cells and normal cells. However, we also found that for samples with more cell types, clustering is not necessarily reduced after imputation, as we many samples cluster around win rate = 0, and the score correlation coefficients between imputed and raw profiles are positive and significant (PCC=0.90, p-value=1.19e-18 for NMI, and PCC=0.78, p-value=7.19e-11 for ARI), which demonstrate that the imputation profile can also preserve most of the biological signals (Figures 4 (e) and (f)). As to the reason why spRefine did not work well in multiple cell-state differentiation, we speculated that the smoothing and denoising effects of imputation on gene expression may have masked some very strong signals that are supposed to be associated with the classification of cell type. Therefore, we recommend checking the predefined number of clusters in the testing datasets before using spRefine as a pre-training framework.

### Predicting survival and disease states for cancer patients from imputed profiles

The imputed gene expression profiles not only provide enriched gene expression information but also help select important features, e.g., marker genes [59] for different conditions. The robust features can also transfer the knowledge from the spatial transcriptomic domain to other omics-type data, for example, predicting the survival information based on the RNA-seq data from The Cancer Genome Atlas (TCGA) [33]. Here we utilized the imputed profiles to select marker genes across the samples from the HEST-1k dataset with different disease states and included the overlapped genes between the marker genes and genes from RNA-seq to predict patient-level survival from the TCGA dataset. Previous research on marker genes has shown that selecting important genes may increase efficiency and improve statistical power [59, 60]. We considered nine cancer types in TCGA, and partitioned the data with RNA-seq and survival information into training, validation, and testing datasets. Here we selected DeepSurv [61] as the baseline method and compared the prediction results between DeepSurv based on marker genes from spRefine and that based on all the genes (Raw) or selected from other sources, including marker genes from scRNA-seq dataset [62] (sc marker) and variable genes [35]. We followed the default setting of DeepSurv and considered three different metrics to evaluate the prediction performance, including concordance td, integrated brier score, and integrated nbll [63].

Details of these metrics can be found in the Methods section, and higher concordance td, as well as lower integrated brier score and integrated nbll indicate better results.

The results for the Raw and spRefine settings for COAD samples are summarized in Figure 5 (a), where spRefine outperformed the Raw setting based on all three metrics, and made significant improvement evaluated by concordance td and integrated nbll. The running time based on spRefine was also 37.2% faster than the Raw mode. Furthermore, we visualize the scores of concordance td based on three different gene lists based on the GBM samples because we can only access the cancer markers from GBM samples at the single-cell resolution in Figure 5 (b). In this figure, the performance based on spRefine is comparable with the Raw setting while outperforming the other two feature selection approaches. This suggests that spRefine offers better marker selections than the other methods. As selecting marker genes might not always improve prediction performance, we plot the prediction performances across nine diseases in Figure 5 (c), choosing marker genes improves prediction in five (COAD, COADREAD, GBM, PRAD, READ) of the nine conditions.

**Fig. 5.**
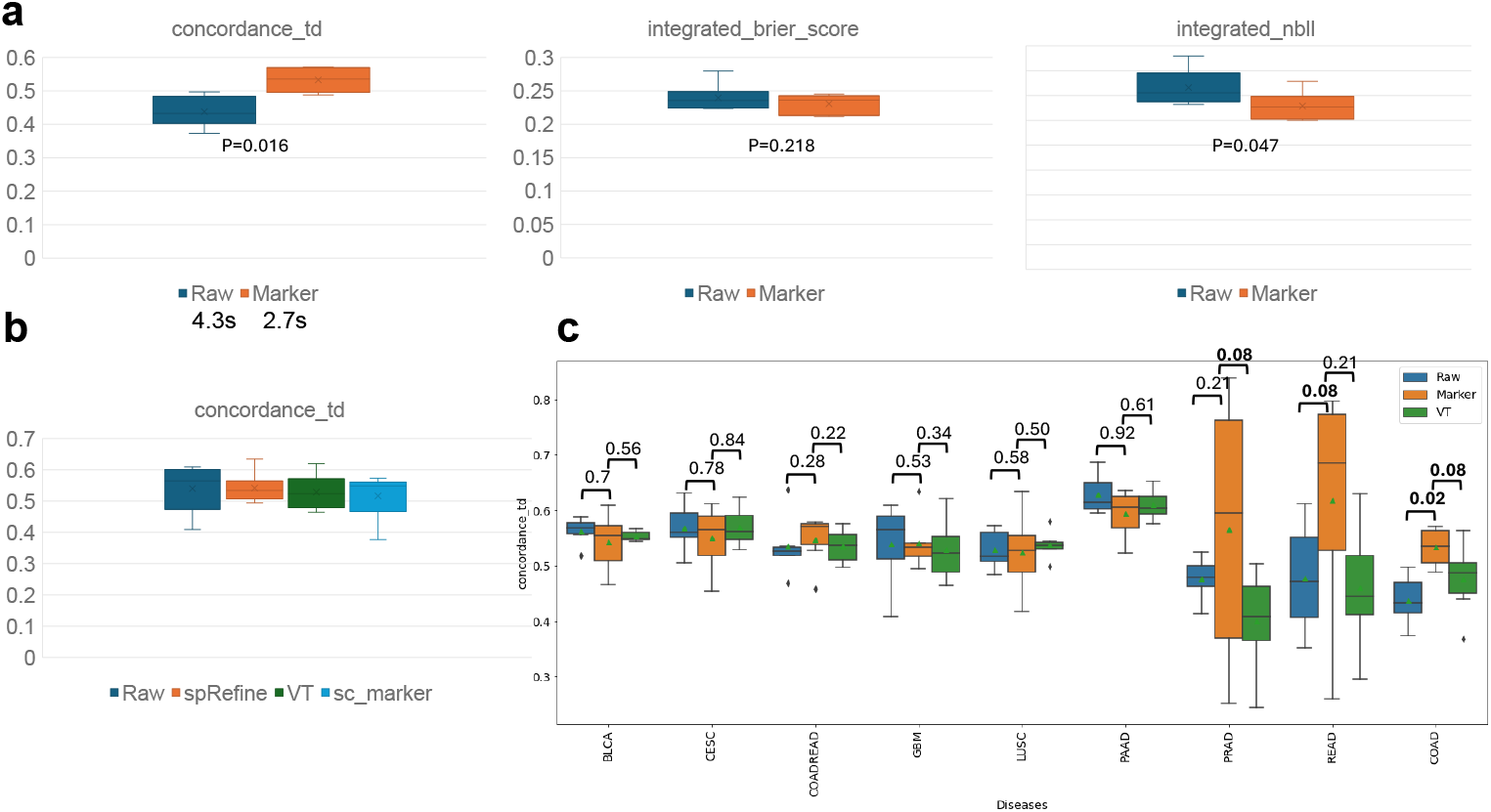
Applications of spRefine for disease modeling. (a) Comparison of survival prediction performances based on three different metrics between raw expression profiles and expression profiles only containing genes selected by spRefine. The cancer type of selected data is COAD. We highlighted the running time comparison and the Wilcoxon Rank-sum test (one-side) p-value (P) in this panel. (b) Comparison of survival prediction performances across marker genes from different sources. (c) Comparison of survival prediction performances across different gene sets based on the cancer types from different cancer types. We annotated the Wilcoxon Rank-sum test (one-side) p-value (P) in this panel, and highlighted the significant result (p-value*<*0.1).

Furthermore, the imputed gene expression profiles can improve the prediction of disease states, leading to a better understanding of diseases. Here we compared classification performance using either the raw expression profiles or imputed expression profiles. We used logistic regression and traditional classification metrics for evaluating [35], and we kept the same parameters for a fair comparison. According to Extended Data Figure 6, using the imputed profiles can improve classification accuracy by achieving higher precision and weighted F1 scores, with similar accuracy.

### Imputing spatial transcriptomics with spRefine can better characterize the spatial ageing clock

Understanding brain ageing can help us reveal the cause of neurodegenerative diseases [13, 64, 65], whose risk increases with aging. The spatial transcriptomic data measured in the brain stratified by age groups provide valuable resources to analyze the effect of spatial context towards ageing and local cell-cell interaction in the ageing process. However, one challenge of building an ageing model is the lack of measured genes, especially for genes associated with ageing effects. The subcellular-level platforms such as MERFISH [66] or STARMAP [8] can only measure expressions from a gene panel and we need to fill the gene expression levels of unmeasured genes. Here we refer to the pipeline of the spatial ageing clock [13], which is a model trained with spatial transcriptomics to predict the age of testing data. We have modified the pipeline by first imputing the gene expression profiles based on spRefine and then estimating the ageing effect by considering the cellular interaction in the spatial context with a graph neural network. The modified pipeline is illustrated in Figure 6 (a). We considered the mouse brain dataset from the original paper on the spatial ageing clock. We first demonstrated the contribution of spRefine by showing the larger number of overlapped genes associated with ageing after imputation (*∼* 1000 genes), shown in Figure 6 (b). The model of the spatial ageing clock has a default gene set used for training and inference, with 100 genes not included in the gene panel, and only two genes not included after imputation. We further evaluated the prediction performances by showing the distribution differences in Figure 6 (c), with the difference statistically significant by the Mann–Whitney U test. Therefore, after imputation with spRefine, the refined spatial ageing clock identified most cells in their correct age group. Our imputed genes can also reflect the differences of cells in the two age groups, shown in Figure 6 (d). Here we visualize the differentially expressed genes (DEGs) identified by the Wilcoxon Rank-sum test between the young (AC124742.1) and old group (Insc), and we can observe specific expression patterns at the spatial level. The earlier expressed genes tend to express in the embryo tissues or early-stage neuron tissue [67], while the later expressed genes tend to express in mature tissues [68].

**Fig. 6.**
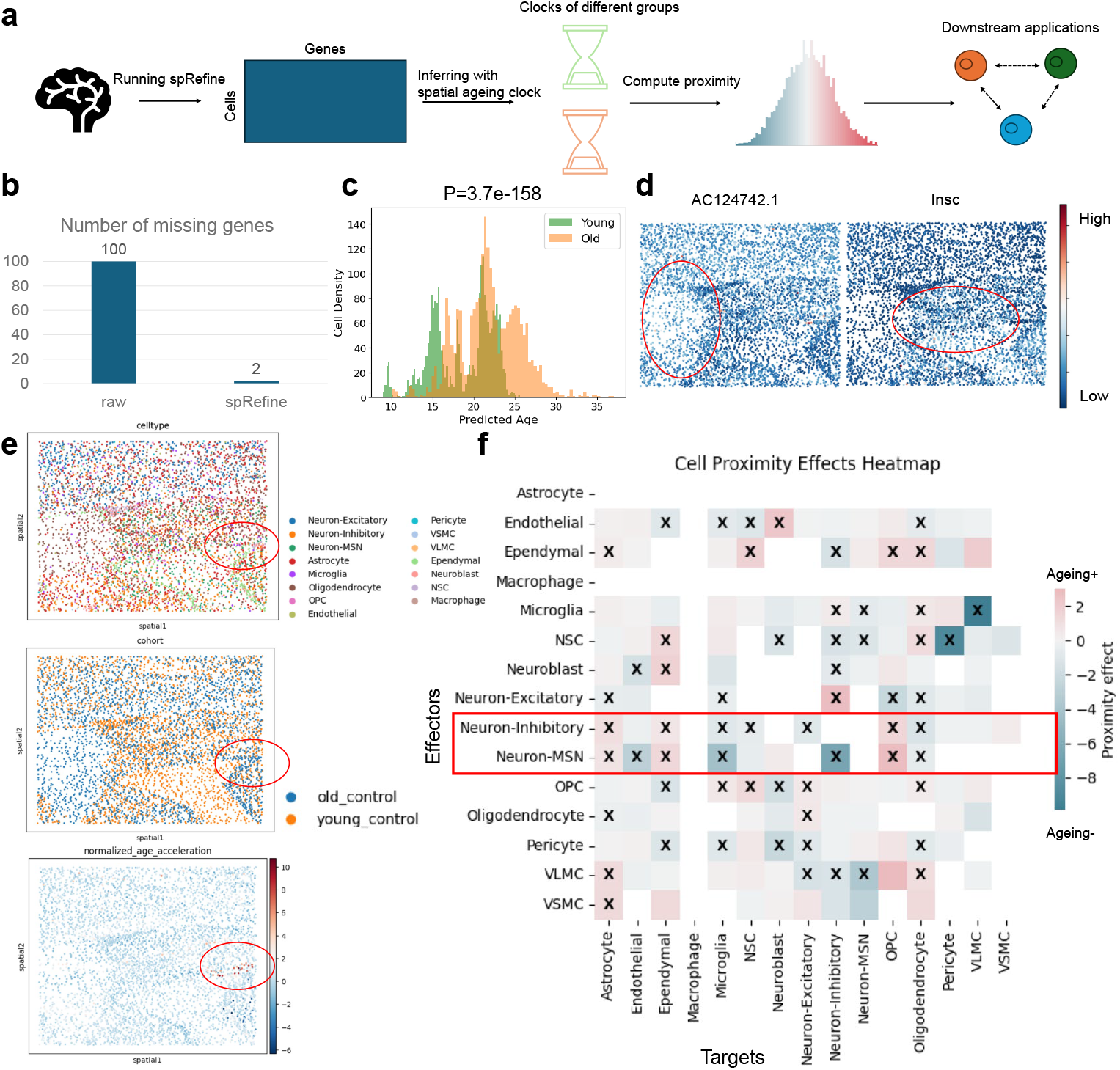
Leveraging contributions of spRefine to understand the biological structure of ageing at the spatial level. (a) The pipeline of ageing effect estimation with spRefine. (b) Comparison of the missing genes between the raw gene set and the imputed gene set. (c) The density of cells predicted with different ages, and the colors represent the measured age group. (d) The spatial-level expression patterns of two selected DEGs from different age groups. Highly expressed regions are highlighted. (e) Visualization of spots colored by cell types, age groups, and normalized age acceleration rate based on spatial location. The area with a strong ageing association was highlighted in a red circle. (f) Heatmap of cell proximity effects by measuring the interactions of different cell types.

In Figure 6 (e), we show the spatial context of the given sample colored by different information, including cell types (upper panel), age groups (middle panel), and normalized ageing acceleration effects (lower panel). Higher acceleration effects mean faster ageing speed rate, which is highlighted in red, which contains cells from both young and old groups, and the cellular composition of this region is also more complex. Therefore, we need to analyze the effects of cell type on senescence by analyzing cellular interactions. Following the ageing effect analysis in the spatial ageing clock pipeline, we calculated and tested the ageing effect of local cell-cell interaction based on the comparison between the nearby cells and distant cells, defined as cell proximity and shown in Figure 6 (f). We used the cross to mark cell-cell interaction with p-value*<* 0.05, where a higher proximity value represents a stronger ageing effect. Compared with the proximity estimation based on the raw expression profiles, the spatial ageing clock based on spRefine can identify more significant cell-cell interactions. Cell types such as Neuron-Inhibitory and Neuron-MSN have the largest number of significant interactions with other cells, which is consistent with recent experimental results for the ageing effect such as neuron function loss in the mouse brain [69, 70]. The third-ranked cell type is NSC, which was reported by the original spatial ageing clock paper as an informative cell type. Moreover, we also discovered several less-explored cell-cell interactions with significant ageing effect, including Endothelial+Ependymal, Ependymal+OPC, Microglia+VLMC, and VLMC+Neuron-Inhibitory. Considering the important roles of these cell types in brain functions, these interactions are worth future investigation.

Finally, we performed external validation based on the scRNA-seq data [65] with mouse brains of different ages, summarized in Figure 7. As shown in Figure 7(a), the selected single-cell dataset has clear cell-type patterns with nearly no batch effect. We also computed the spatial ageing clock information based on this dataset, including predicted age distribution (shown in Figure 7 (b)) and predicted normalized age acceleration effect (shown in Figure 7 (c)). By comparing the estimation results from spatial data and single-cell data, we found that the spatial ageing effect estimation is more precise based on spatial data, which is supported by the clearer age distribution difference based on the imputed spatial data. Furthermore, we computed the cell proximity effects based on the principal components of scRNA-seq data, shown in Figure 7 (d), and after comparing it with effects estimated based on spatial data, we highlighted the cell-cell interactions with the same significance and ageing effect in the same figure. We found overlapped patterns between cell types in these two datasets, such as Neuron cells as effectors and Microglia cells as targets. We visualize the overlapped DEGs based on ages between two datasets, shown in Figure 7 (e) for genes associated with a younger age and Figure 7 (f) for genes associated with a younger age. These selected genes show similar expression patterns in overlapped cell types such as Neuroblast and Oligodendrocyte. Overall, spatial data have higher-resolved cell types and we discovered more cell-cell interactions with experimental support. As a result, the summary based on spatial transcriptomics is more comprehensive, which is also supported by the higher PCCs between predicted and recorded ages estimated from spatial transcriptomics shown in the Figure 2 of [13].

**Fig. 7.**
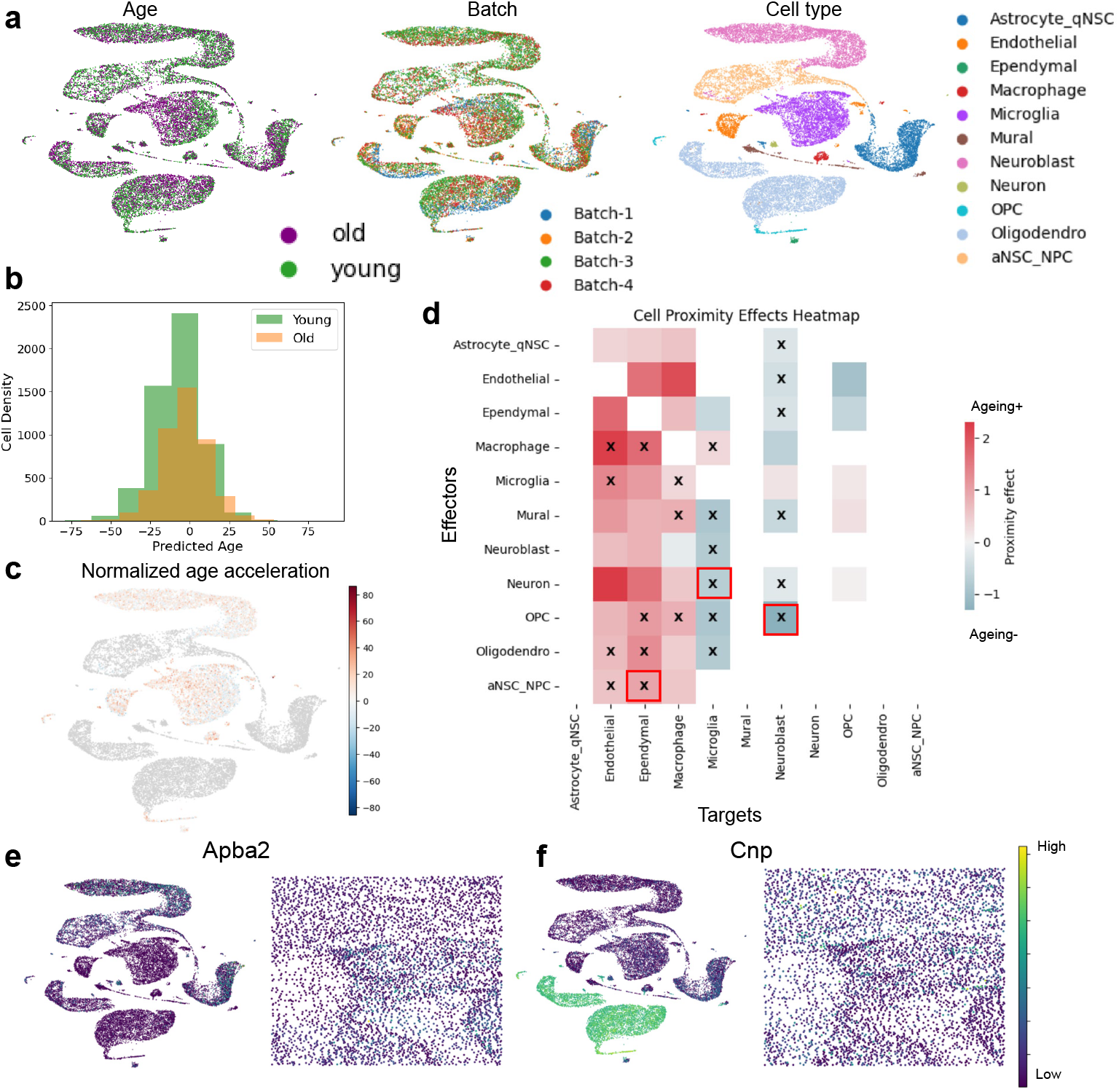
The ageing effect analysis based on scRNA-seq data. (a) The UMAP visualization of measured scRNA-seq data colored by age (left panel), batch label (middle panel), and cell type (right panel). (b) The cell density of predicted cell age colored by observed age information. (c) The UMAP visualization of measured scRNA-seq data colored by normalized age acceleration effect. (d) The cell proximity effect of cell types based on the scRNA-seq data. (e) and (f) represent examples of expression patterns of age-specific marker genes shared by both scRNA-seq data and spatial transcriptomics. Apba2 (e) is a marker from young group and Cnp (f) is a marker from old group. For each gene, the left panel represents UMAPs of scRNA-seq data and the right panel represents spatial location of spatial transcriptomics. Each panel is colored by gene expression levels.

Therefore, the spatial ageing clock estimated from the imputed expression profiles not only preserves the biological signals validated by prior knowledge but also inspires new directions to further investigate the roles of certain cell types in the brain ageing process.

## 3 Discussion

Spatial transcriptomics can provide new insights into understanding biological processes and disease mechanisms in the spatial context. However, there are many challenges in processing spatial transcriptomics, including the imputation of unmeasured genes and the possible denoising of raw gene expression profiles. With imputation and denoising, we can obtain spatial transcriptomics with better coverage and more accurate gene expressions, which can facilitate downstream analyses.

In this manuscript, we have proposed spRefine, a reference-free imputation and denoising pipeline for spatial transcriptomic data analysis, for imputation and denoising. spRefine takes measured spatial transcriptomics as inputs, and utilizes gene embeddings from DNA foundation models to establish gene-gene interactions. We then integrated these two data types to impute unmeasured genes and reduced the noise in the measured genes. Through the analyses of several spatial transcriptomic data, we showed that the results produced by spRefine can better cluster cells and enable more effective downstream analyses.

Given the growing volume of spatial transcriptomic data, there is the potential to collect a large amount of data for pre-training a model to analyze these data. We demonstrated that spRefine can be formatted as a pre-training framework for large-scale spatial transcriptomics collected from different platforms. Our results showed that the imputed expression profiles generated using the pre-trained model can enhance cell classification for some datasets, especially for data with fewer target cell states. Meanwhile, the gene expression profiles obtained after training on large-scale data can also be used for several downstream analyses related to diseases, including the extraction of marker genes to improve survival prediction, and better prediction of sample-level disease state. Therefore, spRefine can serve as a useful pipeline for building foundation models for spatial transcriptomics.

spRefine can also identify cell-cell interactions and spatial ageing effects, which may be highly associated with certain cell states or niches. By imputing gene expression levels, we can obtain a profile with more genes to infer cell-cell interactions in the given sample. Moreover, imputing the ageing-associated genes can identify more ageing proximity cell-cell interactions and thus have a better model to describe the spatial ageing clock. All these novel biological findings demonstrate the potential of the reference-free imputation framework.

Finally, we note that the performance of spRefine may be affected by the choices of models to generate the gene embeddings, and thus a better model can improve the performance of spRefine. In addition, considering that there are only a few publicly available large-scale spatial transcriptomic datasets, while some of them are not well-annotated, collecting and organizing spatial transcriptomic data is also an important research direction. The improvements of both fields may enhance the performance of spRefine to the next level.

## 4 Methods

### Problem definition

In this manuscript, we consider a dataset containing multiple (*m*) spatial transcriptomic profiles, which can be denoted as 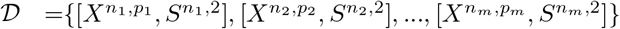, where *X* represents the expression profile and *S* represents the location information. Our target is to utilize the reference gene panel *G*^*q,e*^ with *q* genes and *e* dimensions, where *q >> p*, to impute the unmeasured gene expression in *X* with the final imputed and denoised expression profiles denoted as *X*^*n,q*^. Formally, our target is to learn a function ℳ, which can perform:

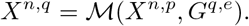

### Model architecture

spRefine considers two components as inputs, including a cell embedding encoder *CE*_*e*_ and a gene embedding encoder *GE*_*e*_. The former takes raw spatial data as input, and the latter takes raw gene embeddings as input. Our gene embeddings are computed based on the pre-trained Enformer [31]. We tested other choices and they did not work as well as the setting based on Enformer. The loss is computed based on the measured genes between the raw input data and imputed outputs, and thus this process can be represented as:

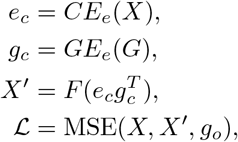

where *g*_*o*_ represents the set of overlapped genes and MSE represents the mean squared error loss function modified for a selected gene set. *F* is a function used to constrain the output format. If we expect to have imputed profiles equal to or larger than 0, then *F* = *Softplus*(); otherwise *F* = *Identity*(). Considering the density of measured gene expression profiles in the selected gene panel design of spatial transcriptomics, we do not model the distribution of outputs in other distributions, but keep on using Normal distribution to compute the loss.

To use spRefine as a pre-training framework, our idea is to impute the missing gene expression profiles proportion to the magnitude of expression levels, which can be defined as a downsampling strategy used in [71, 72]. The model architecture is the same for the imputation mode, while we first mask the input profiles based on a Poisson distribution and fill the masked value based on the imputed results, that is:

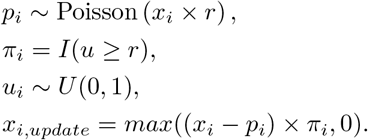

Here *r* = 0.605 represents the dropout rate [71], Poisson represents the Poisson distribution, *I* represents the indicator function, and *U* represents the uniform distribution. We use *x*_*i*_, which represents the raw data from spot *i*, and *x*_*i,update*_ represents the data after downsampling selection, which is the final input for pre-training. Therefore, for the given cell, the final pre-training loss is:

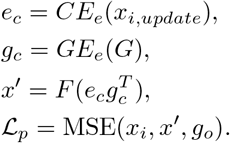

Obviously, this design can impute the missing genes by introducing a set of gene embeddings with unmeasured genes. Moreover, the gene-gene interaction as well as the target of learning a mean value of the measured gene expression profiles can help us reduce the noise level of the given dataset, discussed in [73].

### Model training and hyper-parameter

We utilize PyTorch-lightning [74] to construct and train the model. Our default optimizer is Adam [75], with a learning rate of 1e-4. Our optimal batch size and latent dimensions are tuned for dataset-specific settings, while the batch size is set as large as possible by default, which is shown in Extended Data Figures 7 (a)-(c). We also use the early-stopping method to reduce overfiting and split the original data into training, validation, and testing datasets to evaluate the performance of unsupervised learning.

### Ablation test

When we design spRefine, we consider various ablation tests for the model training stage, including different choices of gene embeddings [76], whether using GNN to encode spot embeddings, whether using variation auto-encoder, whether modeling the decoder output with distribution other than Normal distribution. The results of these settings are shown in Extended Data Figures 2 (a)-(c). We found that making the current model more complicated does not improve its performance, and thus our current design is efficient and optimized.

### Applications

spRefine is capable of various downstream applications after imputation and denoising, including cell-cell communication (CCC) discovery, cell-state stratification, survival prediction, and disease-state prediction.

1. CCC discovery. By using COMMOT [51], we can identify the cell-cell interactions with spatial context and visualize the communication scores for sender genes and receiver genes. We can also visualize the signal flow existing in the given tissue sample, which can help us describe the microenvironment of the disease in a higher resolution.
2. Cell-state stratification. By pre-training spRefine with large-scale spatial transcriptomics, we can generate spot embeddings within the context of diverse disease states. Thus, the provided spot embeddings might work better for identifying disease-associated spots and regions for a given slide. Here we demonstrate that the embeddings from spRefine can be directly used to identify cells with different states and also explain the performance difference affected by the cell-state resolutions.
3. Survival prediction. Traditional methods for survival prediction start from transcriptomic data and train models to predict survival based on the expression profiles of all measured genes. Here we demonstrate that by using the marker genes extracted from the imputed gene expression profiles, we can better predict the survival function of the given samples across different cancer types, and using a smaller number of genes can also improve the model efficiency.
4. Disease-state prediction. We show that using the imputed gene expression profiles from large-scale spatial transcriptomics can help infer the disease state for a whole-slide sample, which further demonstrates the potential of unsupervised learning.
5. Spatial ageing clock construction. We use spRefine to impute the MERFISH sample with ageing information and offer new insights for analyzing the spatial ageing clock model by identifying ageing-associated marker genes, ageing-associated spatial context, and ageing-associated cell-cell interactions. The pipeline we used is the same as the default setting, while the input data are imputed.

### Metrics

Here we discuss the metrics we used to evaluate the performances of spRefine and other baselines in the tasks we performed in this manuscript.

1. For imputation and denoising, we considered NMI, ARI, and ASW as metrics. Details are discussed below:

- Normalized Mutual Information (NMI): We calculate NMI score based on computing the mutual information between the optimal Louvain clusters and the known cell-type labels and then take the normalization. Therefore, NMI *∈* (0, 1) and higher NMI means better performance.
- Adjusted Rand Index (ARI): We calculate ARI score by measuring the agreement between optimal Louvain clusters and cell-type labels. Therefore, ARI *∈* (0, 1) and higher ARI means better performance.
- cell-type Average Silhouette Width (cASW): Here we only consider ASW for cell types. For one cell point, ASW calculates the ratio between the inner cluster distance and the intra cluster distance for this cell. Therefore, higher *ASW*_*cell*_ means better biological information preservation. For *ASW*_*cell*_, we take the normalization, that is:

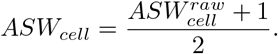
- Avg: This metric represents the average score among these three metrics. All of the metrics are in (0,1) and higher values mean a better model.

2. For batch effect correction, in addition to the three metrics above, we have added more metrics. For metrics covering the evaluation for *S*_*bio*_, including NMI, ARI, cell-type ASW (cASW), and cell-type Local Inverse Simpson’s Index (cLISI). For metrics covering the evaluation for *S*_*batch*_, including bLISI, bASW, KBET, GC, and PCR comparison score.

- Isolated Label Score: Isolated Label computes the batch correction score for the labels within the least number of batches, which offers weights for isolated clusters. Here we utilize the default ASW mode of isolated label score, which means we compute the ASW score for spots from the given label as the final score.
- Local Inverse Simpson’s Index (LISI): This metric is used to evaluate LISI is a metric to evaluate whether datasets are well-mixed under batch labels (*bLISI*) or can be discerned with different cell types *cLISI*. We first compute the k-nearest-neighbor list of one cell and count the number of cells that can be extracted from the neighbors before one label is observed twice.
- batch Average Silhouette Width (bASW): For one cell, ASW calculates the ratio between the inner cluster distance and the intra cluster distance for this cell. The bASW is computed by treating cluster labels as batch labels, and we took the inverse of the computed value to obtain bASW.
- kBET [77]: the kBET algorithm is used to determine if the label composition of the k-nearest-neighbors of a cell is similar to the expected label composition. For the batch label mixture of cells in the same cell types, the proportion of cells from different batches for the neighbors of one cell should match the global level distribution.
- Graph Connectivity (GC): GC measures the connectivity of cells in different cell types. If the batch effect is substantially removed, the connectivity of cells of the same cell type from different batches will have a higher connectivity score based on the k-NN neighbor graph. Therefore, we can compute the GC score for each cell type and take the average.
- PCR Comparison score: PCR is a metric to evaluate the performance of batch effect correction. We calculate the *R*^2^ for a linear regression of the covariate of interest onto each principal component. The variance contribution of the batch effect for all the PCs is based on the sum of the product between the variance of each PC and the *R*^2^ of each PC across all the PCs. Therefore, the score can be represented as:

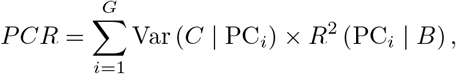

where *G* denotes the number of PCs and *B* denotes the batch information.

For evaluating batch effect correction, all metrics are in (0,1) and higher scores represent better preservation of biological variation or better correction of batch effect. To compute the final score *S*_*final*_, we compute the weighted average: *S*_*final*_ = 0.6*S*_*bio*_+ 0.4*S*_*batch*_. This paper (cite) has demonstrated that the weighted setting will not affect the ranking of models and thus we keep the default setting.

3. Cell-state stratification. Here we directly evaluate the ability of using the embeddings from pre-trained spRefine to identify the cell states based on different samples. Here we compare the clustering metrics (NMI, ARI, and ASW) between the clustering scores from the raw dataset and the imputed dataset.

4. Survival prediction. To evaluate the model performance for survival prediction, we consider three metrics to evaluate the prediction accuracy, including concordance td score, integrated brier score, and integrated nbll score [78].

- concordance td score: The time-dependent concordance index (C-index) is a measure used to evaluate the predictive accuracy of survival models. It quantifies how well the model predicts the ordering of individuals’ event times. Higher score represents better model performance.
- integrated brier score: The integrated Brier score measures the mean squared difference between the observed outcomes and the predicted probabilities at various times. Lower score represents better model performance.
- integrated nbll score: The Negative Binomial Log-Likelihood score measures the probability of the observed outcomes given the model predictions. Lower scores represent better model performance.

5. Disease-state prediction. To evaluate model performance for this task, we utilize well-known metrics for classification performance evaluation, including Accuracy and Weighted F1-score, based on scikit-learn [35]. Higher score represents better model performance.

### Baselines

Here we consider different baseline models for different tasks. The methods are ranked in alphabet (A-Z) order.

#### 1. Imputation and denoising

- ENVI [20] is a model based on conditional auto-encoder. It learns the embeddings of scRNA-seq data and multiplexed spatial transcriptomics simultaneously and decodes the embeddings to expression space for imputation. However, ENVI meets OOM issues in our benchmarking process.
- gimVI [19] also models the gene expression from these two modalities into a joint latent space and uses variational inference to generate the output distribution and impute the expression levels of missing genes. However, gimVI does not consider the neighborhood relation in the spatial data and its implementation is not efficient, as shown in our results. Moreover, gimVI meets OOM errors in our benchmarking process.
- SpaGE [17] is a model based on dimension reduction and regression. It firstly reduces the high dimensions of the input data, and in the joint low-dimensional space, it trains a regression model to impute the value of missing genes. However, SpaGE is not efficient for Xenium-based data with moderate performance.
- Tangram [11] is a model based on optimal matching. Tangram learns the best match relation between scRNA-seq data and spatial data by learning a mapping function and then performs the imputation based on minimizing the loss of such mapping function. However, Tangram is not scalable and the batch version of Tangram does not consider the difference between local optimal solutions and global optimal solutions.
- TransImp [37] is a model based on regression and spatial information regularization. It also relies on dimension reduction for the first step, and in the regression step, it considers both minimizing the loss between predicted data and ground truth data and minimizing the difference of spatial information. However, TransImp meets OOM errors in our benchmarking process and its imputation results lead to poor performance for some downstream applications, for example, RNA velocity inference.
- VISTA [22] models the gene expression profiles based on a coupled graph variational auto encoder. VISTA learns the representations of spots and cells based on encoder and then reduce the distance between two different domains and impute the gene expression profiles measured in the spatial context based on a decoder. VISTA is capable of performing various downstream applications, including CCC discovery, spatial RNA velocity inference, and spatial perturbation simulation.

#### 2. Batch effect correction

- DeepImpute [44] uses a denoising auto-encoder to impute the single-cell transcriptomics. It only considers filling zero-expressed genes in cells.
- MAGIC [43] uses a graph-diffusion map approach to impute the single-cell transcriptomics. It considers imputing both unmeasured gene expression levels as well as enhancing known gene expression levels.
- SEDR [12] utilizes a graph auto encoder to encode the gene expression of spots into a latent space and then reconstruct the masked gene expression profiles to perform unsupervised training. It then utilizes embeddings to enhance other downstream applications, including batch effect correction and clustering.
- STAGATE [42] considers a cell-type-level graph auto encoder as well as a spatial-level graph auto encoder to learn the representation with a weighted-sum approach. It also takes the representation as inputs and decodes them into expression profiles, which have been denoised after training. This method can also identify spatial domains and help extract 3D spatial architecture.

#### 3. Cell-state stratification

- Raw expression profiles. Here we take the gene expression profiles without imputing and denoising as the baseline.

#### 4. Survival prediction

- All gene set. Here we first consider using all of the genes to predict the survival function of the given sample.
- Features from RNA-seq. Here we utilize variance threshold approach to select marker genes from the training dataset, and then test the model performance.
- Marker gene from scRNA-seq. Here we collect the disease marker genes from 3CA database [62] of diseases and select these markers for testing

#### 5. Disease-state prediction

- Raw expression profiles. Here we take the gene expression profiles without imputing and denoising as the baseline.

#### 6. Spatial ageing clock construction

- Raw expression profiles. Here we take the gene expression profiles without imputing and denoising as the baseline.

## Datasets

Details of used data in this paper are summarized in Supplementary File 1.

## Code availability and reproductivity

Our computation resources are based on Yale High-performance Computing Center (YCRC). For data pre-processing, we reply on one CPU with up to 800 GB. For model training, we utilize one NVIDIA A100 (H100) GPU with up to 100 GB RAM.

The codes of this project can be found in https://github.com/HelloWorldLTY/sprefine. The license is MIT license.

## 5. Ethics and Inclusion

Although spRefine is not biased to gender, races, and other factors, the users are solely responsible for the content they generate with models in spRefine, and there are no mechanisms in place for addressing harmful, unfaithful, biased, and toxic content disclosure. Any modifications of the models should be released under different version numbers to keep track of the original models related to this manuscript.

The target of current spRefine only serves for academic research. The users cannot use it for other purposes. Finally, we are not responsible for any effects of the use of the model.

## 6. Acknowledgments

We thank Dr. Zhang Le for suggesting methods to analyze ageing effect in the brain region. This project is partially supported by NHGRI K99HG013429 (Dr. Tinyi Chu), NIH U24 HG012108 and U01 HG013840 (Dr. Hongyu Zhao), and NSF IIS Div Of Information & Intelligent Systems 2403317 (Dr. Rex Ying).

## 7. Author contributions

T.L. proposed the study. T.L., T.H., W.J., and T.C. designed the model. T.L. ran all the experiments. R.Y. provided the computation resources. T.L., T.H., W.J., R.Y. and H.Z. wrote the manuscript. H.Z. supervised this project.

## A Supplementary figures

**Extended Data Figure 1.**
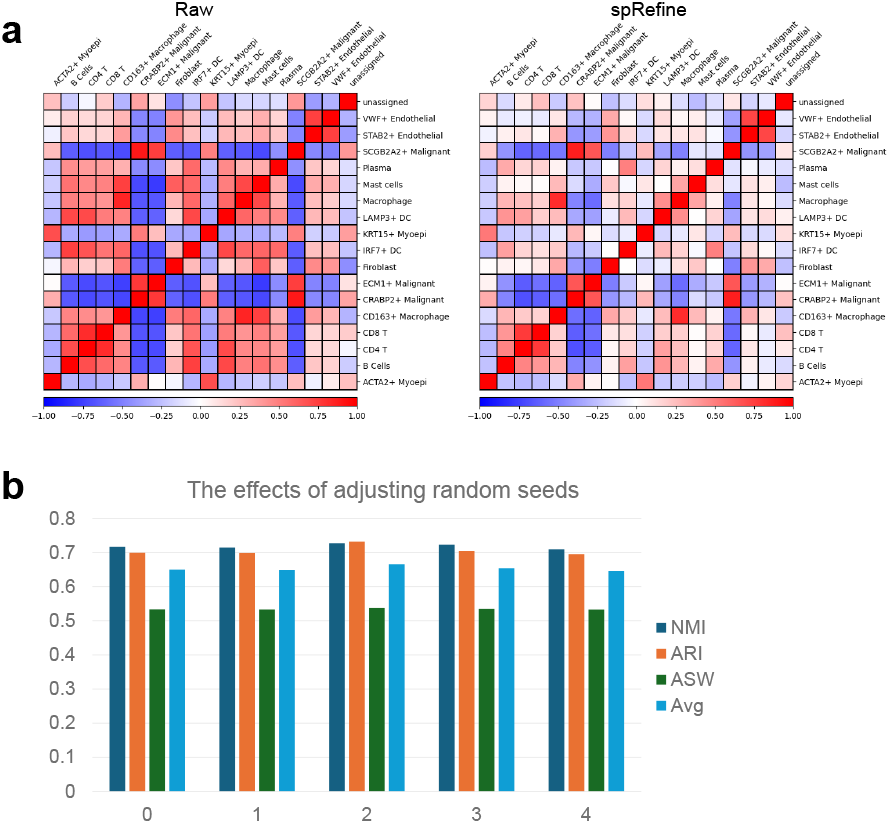
Comparison of cell-type similarity and performance under different random seeds. The dataset analyzed in this figure is xenium breast. (a) Cell-type-level similarity computed based on gene expression levels. The left panel represents the similarity computed based on the raw profile, while the right panel represents the similarity computed based on the imputed profile. (b) Clustering performance of spRefine across different random seeds.

**Extended Data Figure 2.**
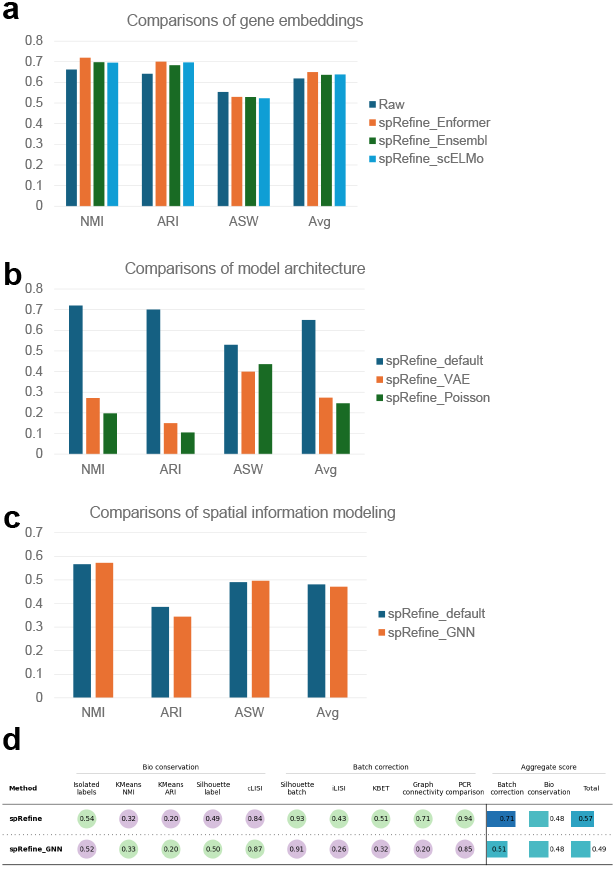
Ablation tests of spRefine. (a) Clustering performance of spRefine with gene embeddings from different sources based on the xenium breast dataset. (b) Clustering performance of spRefine with different model architectures based on the xenium breast dataset. (c) Clustering performances of spRefine with spatial modeling approaches based on the xenium brain dataset. (d) Comparison of batch effect correction performances between spRefine with and without GNN encoder.

**Extended Data Figure 3.**
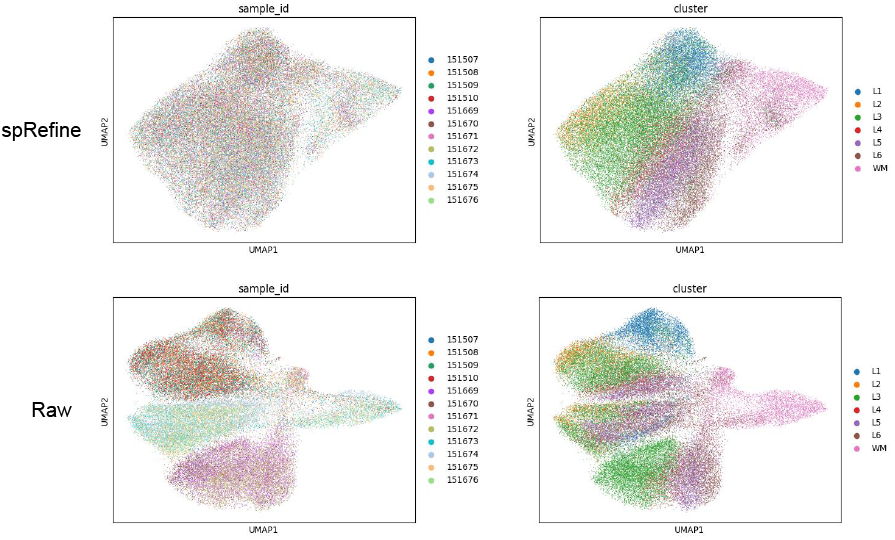
UMAP visualization of gene expression profiles before and after denoising+batch effect correction. These figures are colored by batch labels (sample id) or cell types (cluster).

**Extended Data Figure 4.**
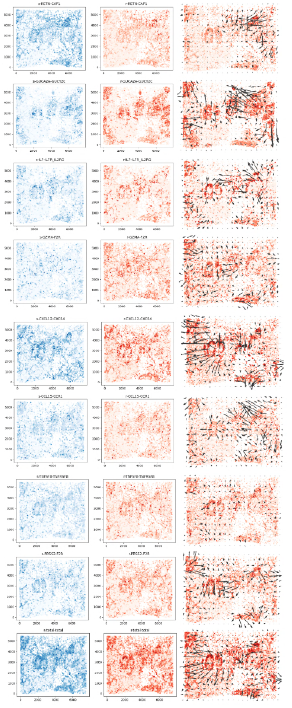
The first part of measured CCCs. Each line contains three panels, including sender strength, receiver strength, and signal directions.

**Extended Data Figure 5.**
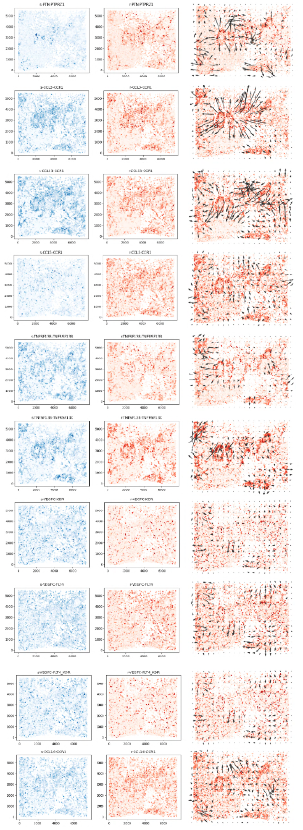
The second part of measured CCCs. Each line contains three panels, including sender strength, receiver strength, and signal directions.

**Extended Data Figure 6.**
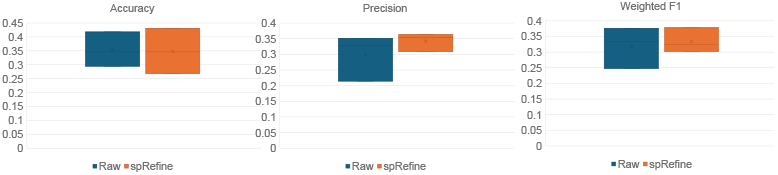
Comparison of disease-state prediction performances between the raw and imputed profiles by spRefine. Here we selected 10% of the cells in the original data to implement the training.

**Extended Data Figure 7.**
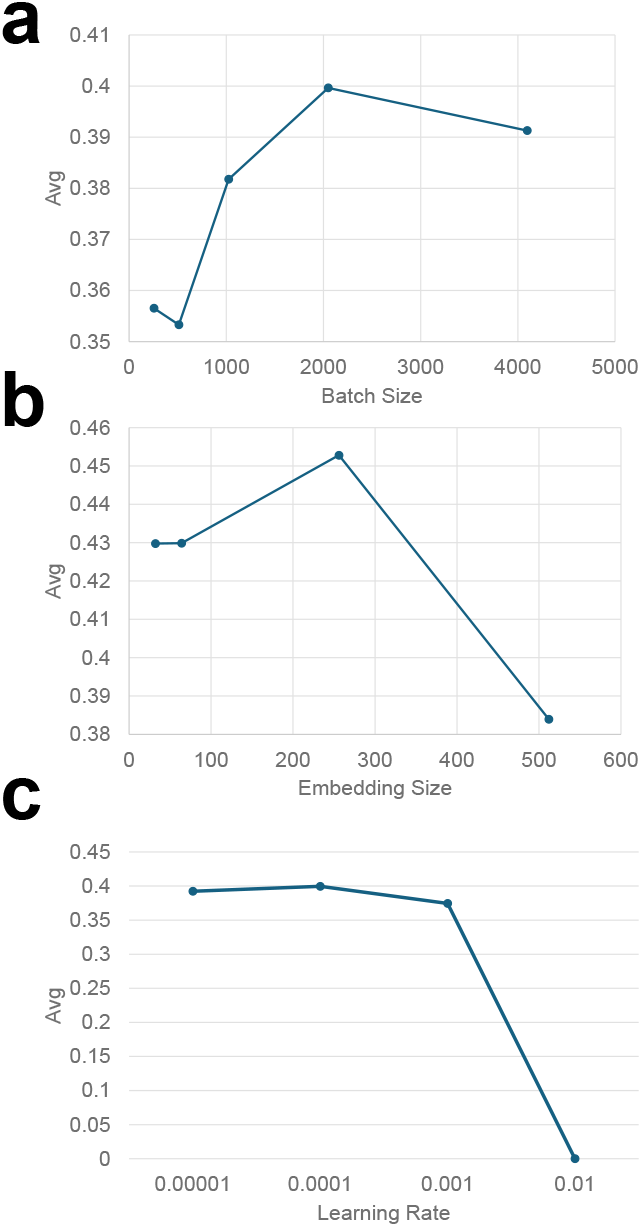
The comparisons of clustering performances based on different hyper parameters. (a) Model performances under different batch sizes. (b) Model performances under different embedding sizes. (c) Model performances under different learning rates.

