## Supplementary figures, supplementary files 1 for "spRefine Denoises and Imputes Spatial Transcriptomics with a Reference-Free Framework Powered by Genomic Language Model": Appendix.pdf

### A Supplementary figures

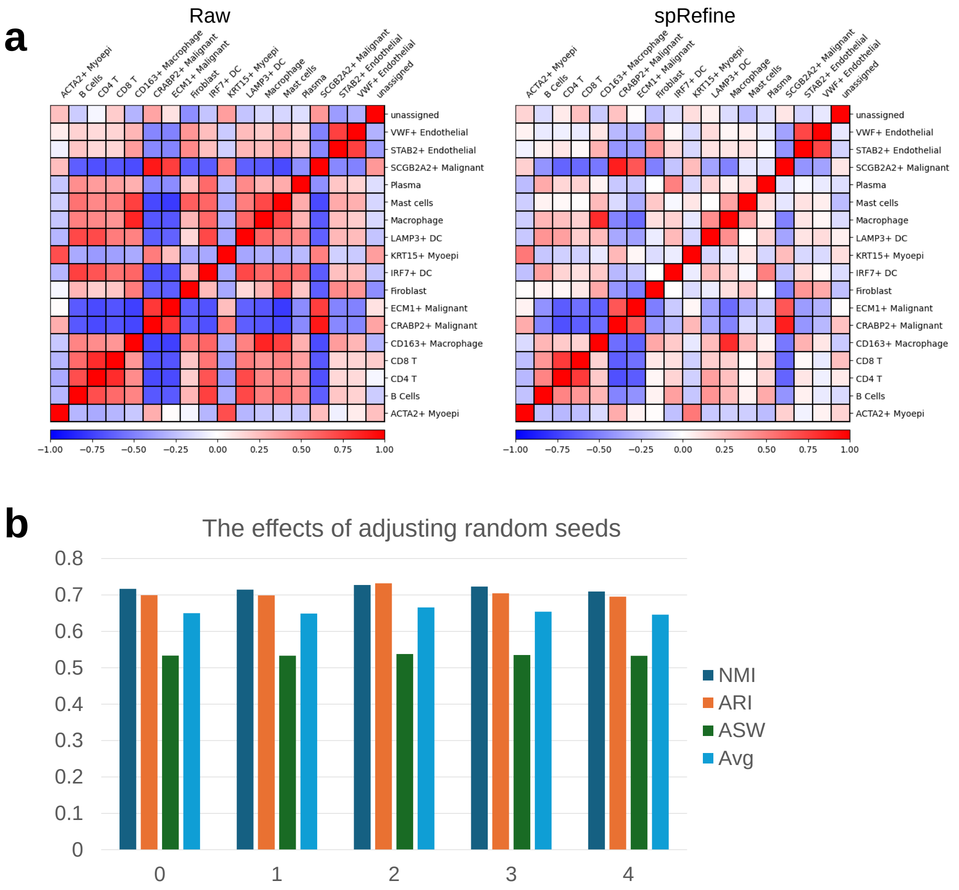


**Extended Data Fig. 1** Comparison of cell-type similarity and performance under different random seeds. The dataset analyzed in this figure is xenium_breast. (a) Cell-type-level similarity computed based on gene expression levels. The left panel represents the similarity computed based on the raw profile, while the right panel represents the similarity computed based on the imputed profile. (b) Clustering performance of spRefine across different random seeds.

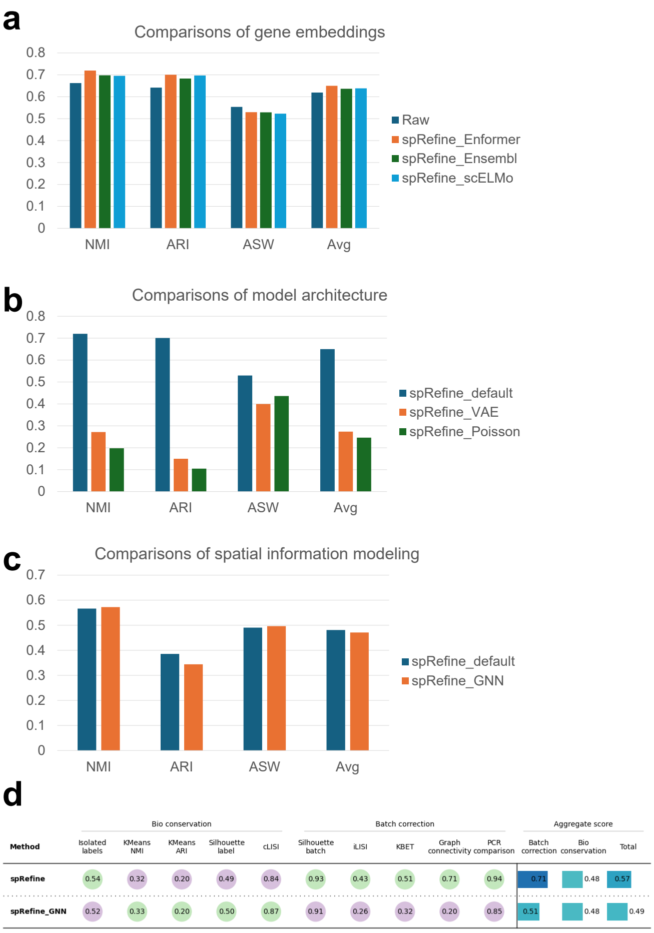


**Extended Data Fig. 2** Ablation tests of spRefine. (a) Clustering performance of spRefine with gene embeddings from different sources based on the xenium_breast dataset. (b) Clustering performance of spRefine with different model architectures based on the xenium_breast dataset. (c) Clustering performances of spRefine with spatial modeling approaches based on the xenium_brain dataset. (d) Comparison of batch effect correction performances between spRefine with and without GNN encoder.

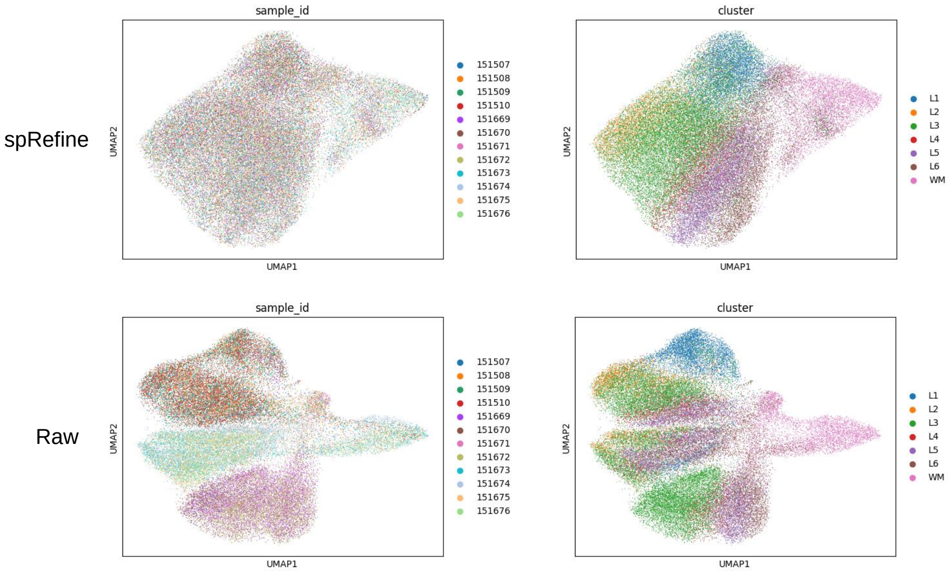


**Extended Data Fig. 3** UMAP visualization of gene expression profiles before and after denoising+batch effect correction. These figures are colored by batch labels (sample_id) or cell types (cluster).

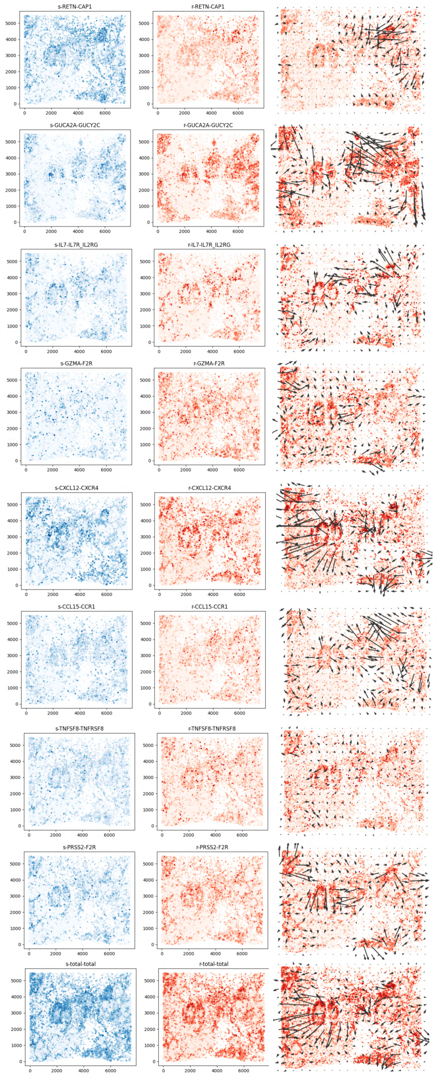


**Extended Data Fig. 4** The first part of measured CCCs. Each line contains three panels, including sender strength, receiver strength, and signal directions.

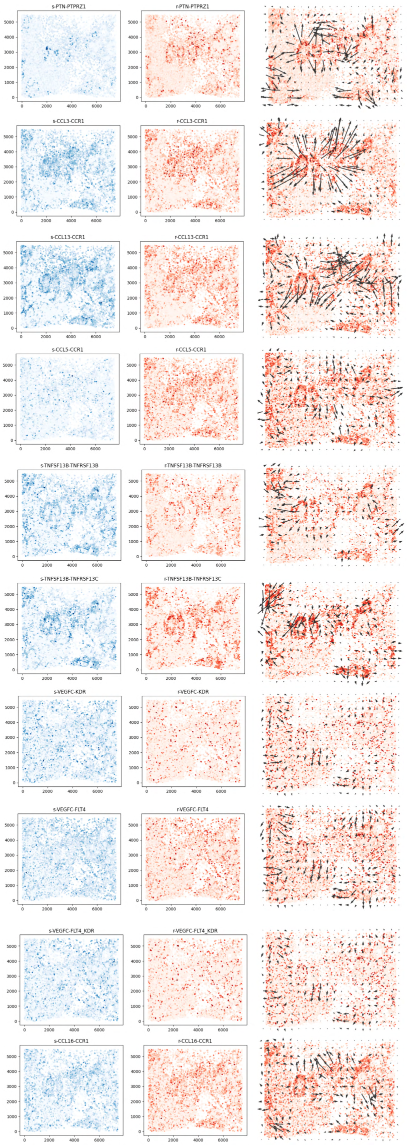


**Extended Data Fig. 5** The second part of measured CCCs. Each line contains three panels, including sender strength, receiver strength, and signal directions.

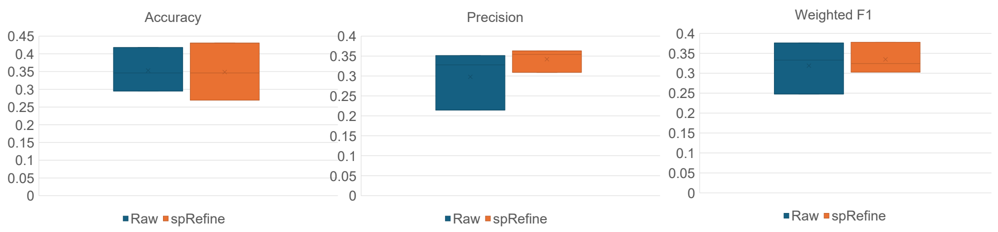


**Extended Data Fig. 6** Comparison of disease-state prediction performances between the raw and imputed profiles by spRefine. Here we selected 10% of the cells in the original data to implement the training.

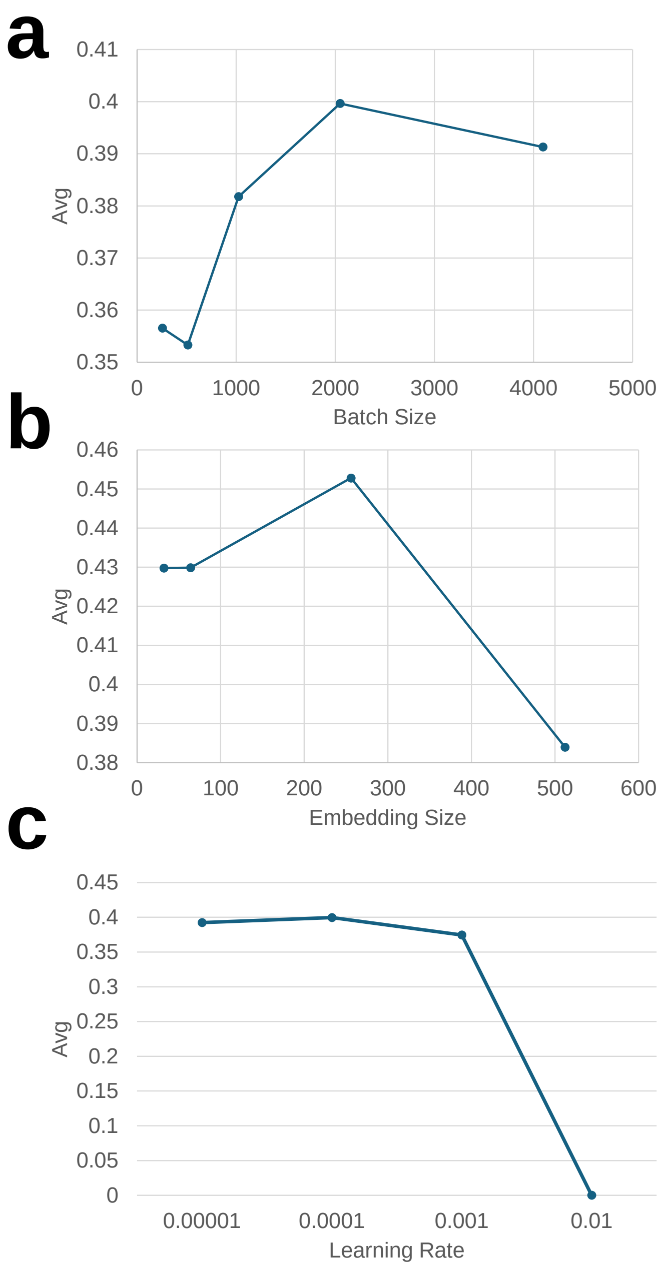


**Extended Data Fig. 7** The comparisons of clustering performances based on different hyper parameters. (a) Model performances under different batch sizes. (b) Model performances under different embedding sizes. (c) Model performances under different learning rates.
